# The role of cuticular hydrocarbons in intraspecific aggression in the invasive ant *Cardiocondyla obscurior*

**DOI:** 10.1101/2023.02.01.526643

**Authors:** Maja Drakula, Jan Buellesbach, Lukas Schrader

## Abstract

Cuticular hydrocarbons (CHCs) are important cues for nestmate discrimination and intraspecific aggression in ants. In invasive ants, diminished CHC profile diversity is suspected to contribute to the ecological and evolutionary success of populations by reducing intraspecific aggression between colonies. The ant *Cardiocondyla obscurior* has successfully colonized habitats around the world, reaching high local population densities. However, despite being invasive, colonies still react aggressively against each other, especially in interactions with non-nestmate alate queens. Here, we study whether CHCs are relevant for antagonistic interactions in this species, by combining behavioral experiments with gas-chromatography coupled with mass spectrometry (GC-MS). We show that queen and worker CHC profiles show pronounced quantitative as well as qualitative differences, that queens with depleted CHC profiles receive virtually no aggression from non-nestmates, and that aggression levels are positively correlated with the naturally occurring CHC profile differences between colonies. These findings provide first empirical evidence for a role of CHCs and chemical diversity in antagonistic behaviors against foreign queens in this species. They further suggest that invasive populations of *C. obscurior* are multicolonial and polydomous.

## Introduction

Communication is a key element in eusocial systems (Holldobler & Wilson 2009). Social insects have evolved sophisticated communication systems, predominantly mediated by chemical cues (Holldobler & Wilson 2009). These cues are particularly fundamental for kin recognition, as they allow for the identification of nestmates and reproductive members of the colony. Therefore, chemical communication requires accurate and fine-tuned coordination in eusocial systems (Holldobler & Wilson 2009, Sturgis & Gordon 2012).

Hydrocarbons, which are secreted onto the insect cuticle along with alcohols, sterols, and other compounds (Millar 2010), are important chemical signals in insect communication (Martin & Drijfhout 2009, Kidokoro-Kobayashi & al. 2012). In ants, colonies commonly carry a distinct colony odor (Sturgis & Gordon 2012, Sprenger, L. and Menzel 2020) most commonly defined by the composition of straight-chain, methyl-branched- and unsaturated cuticular hydrocarbons (CHCs) (Martin & Drijfhout 2009, Sturgis & Gordon 2012). This unifying colony odor can be sustained through trophallaxis (Holldobler & Wilson 2009), grooming or direct body contact (Lahav & al. 1999) between nestmates and varies depending on genotype, environmental conditions, as well as the physiological and nutritional state of individual ants (Crozier & Dix 1979, Martin & Drijfhout 2009, Gruber & al. 2012, Sturgis & Gordon 2012). The colony specific CHC blends are commonly considered crucial for robust discrimination between nestmates and non-nestmates in most ant species (Lahav & al. 1999, Sprenger, L. and Menzel 2020). In populations of invasive ants, the depletion of genetic diversity through genetic bottlenecks is suspected to reduce differences in colony odors between colonies and impede nestmate recognition (Tsutsui & al. 2003, HelanterÄ & al. 2009).

In consequence, intraspecific aggression is often low or absent in populations of invasive ants, allowing the emergence of large networks spanning hundreds of nests that function as a single supercolony (Holway & al. 2002, Fournier & al. 2009, HelanterÄ & al. 2009, Eyer & al. 2018). This unicolonial structure of invasive populations is considered key for the ecological success of many introduced ant species, as it reduces the costs of intraspecific conflicts thus allowing for extreme population densities (Tsutsui & al. 2000, Holway & al. 2002). However, unicoloniality is apparently not a necessary condition for the persistence of invasive populations of ants. For instance, invasive colonies of the ant *Brachymyrmex patagonicus* (Mayr, 1868) engage in antagonistic interactions, resulting in a multicolonial rather than a unicolonial population structure (Eyer & al. 2021).

Similarly, in invasive populations of the heart-node ant *Cardiocondyla obscurior* (Wheeler, 1929), colony boundaries do exist as workers successfully discriminate against non-nestmates. However, foreign workers have been observed to be eventually integrated into a colony after short periods of initial aggression (Heinze & al. 2006). In contrast, foreign alate queens are met with substantially more aggression (Schrader & al. 2014). The mechanisms behind these antagonistic interactions have not been studied so far.

Populations of *C. obscurior* can be found in diverse habitats across temperate, subtropical, and tropical regions around the world, often reaching high local densities (Heinze & al. 2006, Heinze 2017, Errbii & al. 2021). Alate queens in this polygynous species mate with males inside their own nest before founding a new colony by budding (Heinze & al. 2006, Suefuji & al. 2008, Heinze 2017), which creates a dense network of closely related colonies within close proximity. This mode of colony propagation, short generation times, and genomic traits (Schrader & al. 2014, Errbii & al. 2021) likely contribute to the success of *C. obscurior* outside its native range (Heinze & al. 2006, Heinze 2017). However, to what extent invasive populations of this species can be considered unicolonial or multicolonial remains unclear, as empirical data on population structure, colony odor diversity, and antagonistic interactions are still very scarce. To gain further insights into the structure of invasive populations of *C. obscurior*, we studied colonies sampled from a dense invasive population in Bahia, Brazil to address the following questions: (1) Are CHC profiles different between alate queens and workers and how variable are these profiles in a population?, (2) are CHCs vital cues for worker discrimination against non-nestmate queens in this species?, and (3) are CHC profile differences between colonies sufficient to explain aggressive interactions? By combining behavioral assays with CHC profiling based on gas-chromatography coupled with mass spectrometry (GC-MS), we demonstrate (1) that queens and workers have qualitatively and quantitatively distinct CHC profiles (2) that CHCs are key cues for nestmate recognition and antagonistic responses, and (3) that CHC profile divergence between colonies are sufficient to explain aggression.

## Materials and methods

### Ant collection and rearing

Ten colonies of *Cardiocondyla obscurior* collected from two different locations in Bahia, Brazil in 2018 (Una and Itabuna, five colonies each) were used for the aggression assays and CHC analysis (Supplementary Table **S1**). The colonies were kept in artificial plastic nests (10 cm), half filled with plaster and a smaller chambered nest (2 cm x 6 cm) covered with red plastic foil at a constant temperature of 22-26 °C, a 12 h light/dark cycle, 75% humidity. They were fed three times a week with legs of cockroaches, a water-soaked sponge and paper soaked with honey (Blütenhonig, dm-Drogerie Markt GmbH & Co. KG, Karlsruhe, Germany).

### Chemical analysis

For CHC analysis, ten respective individuals of workers and alate queens from each of the ten colonies were pooled and extracted in a 1.5 ml glass vial (Agilent Technologies, Waldbronn, Germany) with 100 μl of MS pure *n-*hexane (Merck KGaA, Darmstadt, Germany). The vials were then put on an orbital shaker (KS 130 basic, IKA^®^-Werke GmbH & Co. KG, Staufen, Germany) for 10 min and 240 rpm, after which the generated surface extracts were transferred into 250 μl micro-volume inserts (Agilent Technologies, Waldbronn, Germany) and dried under a constant stream of carbon dioxide. The sample was resuspended in 5 μl of hexane containing 7.5 ng/μl of dodecane (*n*-C12) as an internal Standard. 5 μl of the extract was then injected using a PAL RSI 120 (CTC Analytics AG, Switzerland) automatic sampler in split/splitless mode into a gas chromatograph (7890B, Agilent Technologies, Waldbronn, Germany) coupled with a Triple Quadrupole mass selective detector (7010B, Agilent Technologies, Waldbronn, Germany, Quad: 150 °C, Source: 230 °C) equipped with a DB-5MS UI capillary column (Agilent Technologies, Waldbronn, Germany, length, 30 m; ID: 0.25 mm; film thickness: 0.25 μm). Helium was used as carrier gas at a constant flow rate of 1.5 ml per min. The temperature program started with 80 °C held for 1 min before increasing to 300 °C at a rate of 5 °C min^-1^, and after a hold time of 5 min to 325 °C at a rate of 5 °C min^-1^, where it ends with a hold time of 5 min.

Chromatograms were first visually inspected using the MassHunter Workstation Qualitative Analysis Software (B.08.00 SP1, Agilent Technologies, California, USA). CHC compounds were identified based on their diagnostic ions and retention indices (RI) (Carlson & al. 1998). The profiles were then compared with consecutive CHC peak detection, integration, and quantification with MassHunter Workstation Quantitative Analysis Software (B.09.00, Agilent Technologies, California, USA).

### Aggression assays

To test to what extent CHC differences in queen profiles correlate with worker aggression, the aggressive response of workers towards untreated queens (“alive”), freeze-killed queens (“freeze-killed”), and freeze-killed queens with depleted CHC profiles (“hexane washed”) were quantified and compared. For CHC depletion, pools of alate queens for each colony were put in a glass vial (1.5 ml) containing 100 μl *n*-hexane for 24 hours. After extracting the hexane, fresh hexane was added and this process was repeated three times, before drying off the hexane under carbon dioxide. To assess the effect of the different treatments on CHC profiles, we characterized CHC profiles of pools of 10 alate queens for each treatment group (alive, freeze-killed, hexane washed) as described above. Compared to untreated individuals, hexane washing removed 91.6 % of CHCs. The CHC profiles of freeze-killed individuals did not differ from untreated individuals.

The behavioral assays were conducted with individuals from ten colonies from two different sites (Una and Itabuna, Brazil). We setup ten experimental colonies consisting of 20 workers and ten brood items in a small nest chamber (6 cm) and a single narrow exit. All colonies were setup on the same day. Colony size was kept constant throughout the experiments by addition of workers and brood from the respective source colony.

We collected 270 alate queens from the ten source colonies (i.e., 27 per colony) for our behavioral assays, designating nine individuals per colony to each of the three treatment groups (i.e., 90 individuals per treatment group). Alive alate queens used in aggression assays were collected from their colony at the day of the experiment and separated into petri dishes (3 cm diameter). The hexane washed and freeze-killed individuals were prepared at the beginning of the experiments and stored at -20°C until further use. For observation during the aggression assays, experimental nests were placed under a Keyence digital microscope (VHX-900F, Keyence, Osaka, Japan), combined with a free-angle observation system base unit (VHX-S90BE, Keyence, Osaka, Japan), and a free-angle observation system lens holder (VHX-S90F, Keyence, Osaka, Japan) with a Z20, X20 resolution. Alive alate queens were introduced to the experiment in 1 ml pipette tips to avoid stress by handling them with tweezers. Freeze-killed individuals were placed at the nest entrance and lightly pushed into the nest cavity of the experimental colonies with fine forceps.

The observer was blind to the treatment of the queens and the recipient/intruder colony.

To quantify aggressive interactions, the behavior of workers towards the introduced alate queens (alive, hexane washed and freeze-killed) was observed for five minutes, recording the highest intensity of aggression expressed by workers using the following ordinal scale: (1) Antennal contact: Antennating the trespassing foreign queen, (2) Mandible flare: Facing the trespassing foreign queen with widely open mandibles, (3) Biting: Biting the trespassing foreign queen in any body part, (4) Pulling: Biting and subsequently pulling the trespassing foreign queen, (5) Stinging: Stinging the trespassing foreign queen.

### Statistical Analysis

All statistical analyses were done in R (R Core Team 2022). To test whether behavioral responses of workers differed significantly between treatments, we fitted an ordinal logistic regression model (aggression∼treatment) implemented in the polr function of the R library MASS (Venables & Ripley 2002, Hothorn & al. 2008) and applied Tukey’s post-hoc test as implemented in the glht function included in the masscomp library (Hothorn & al. 2008). Furthermore, as a quantitative measure of chemical distance in CHC profiles between colony pairs, we conducted a principal component analysis of the CHC colony profiles and calculated Manhattan distances across the first two principal components for all pairs. We then tested for an effect of CHC distance on aggression, using an ordinal logistic regression model with CHC distance as a continuous variable (aggression∼treatment * CHC distance), again using the polr function of the MASS package.

## Results

To explore the role of CHCs in nestmate recognition and intraspecific aggression in *Cardiocondyla obscurior*, we first characterized CHC profiles and their variation for alate queens and workers from ten colonies sampled from an invasive Brazilian population. We identified 71 individual CHC compounds in queens and 61 in workers (**Table 1**). Ten compounds (five unsaturated alkenes and five alkadienes) were exclusive to alate queens (Supplementary Figure **S1**). We did not find any compounds exclusive to workers.

**Tab. 1.**
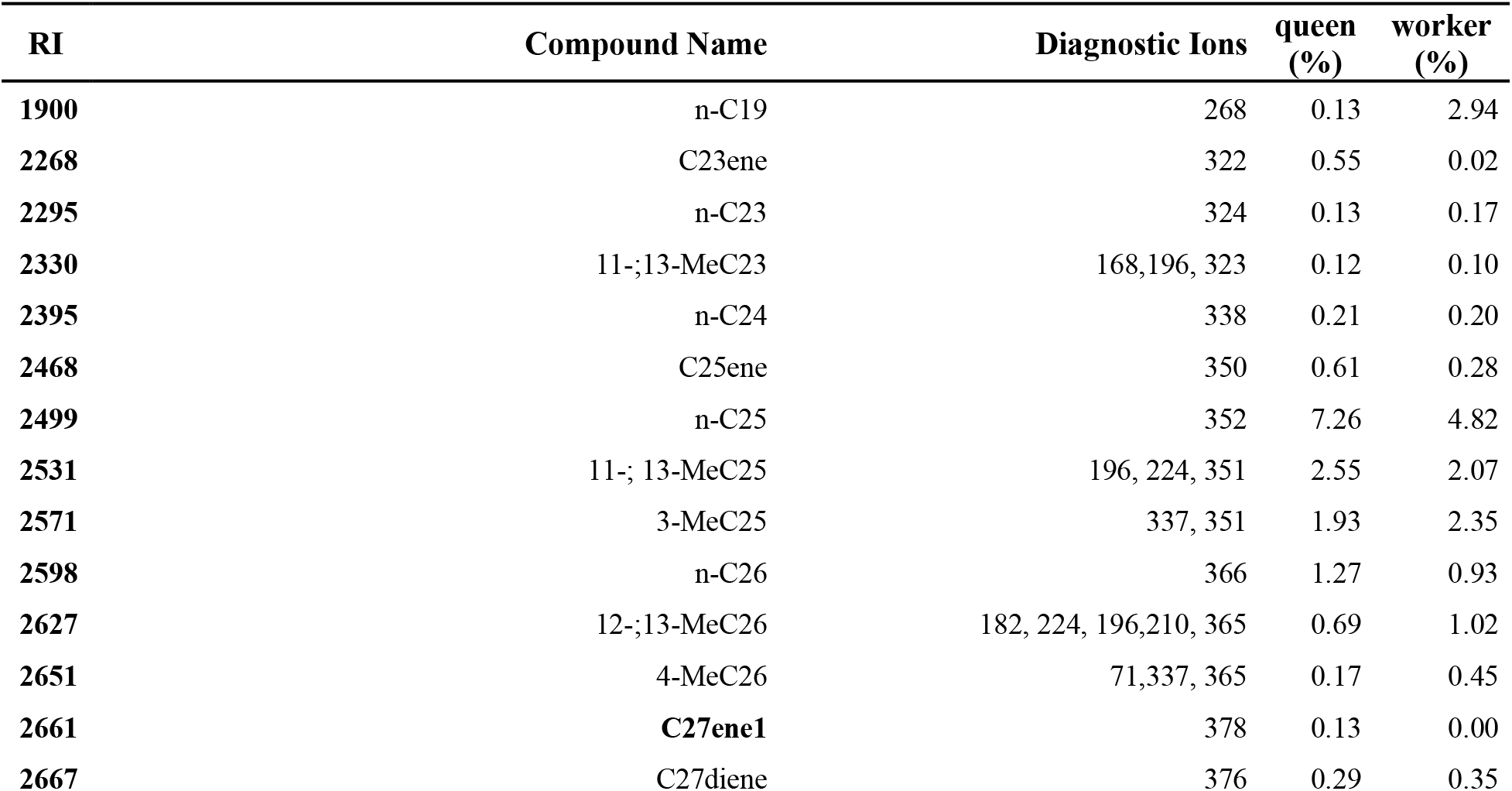
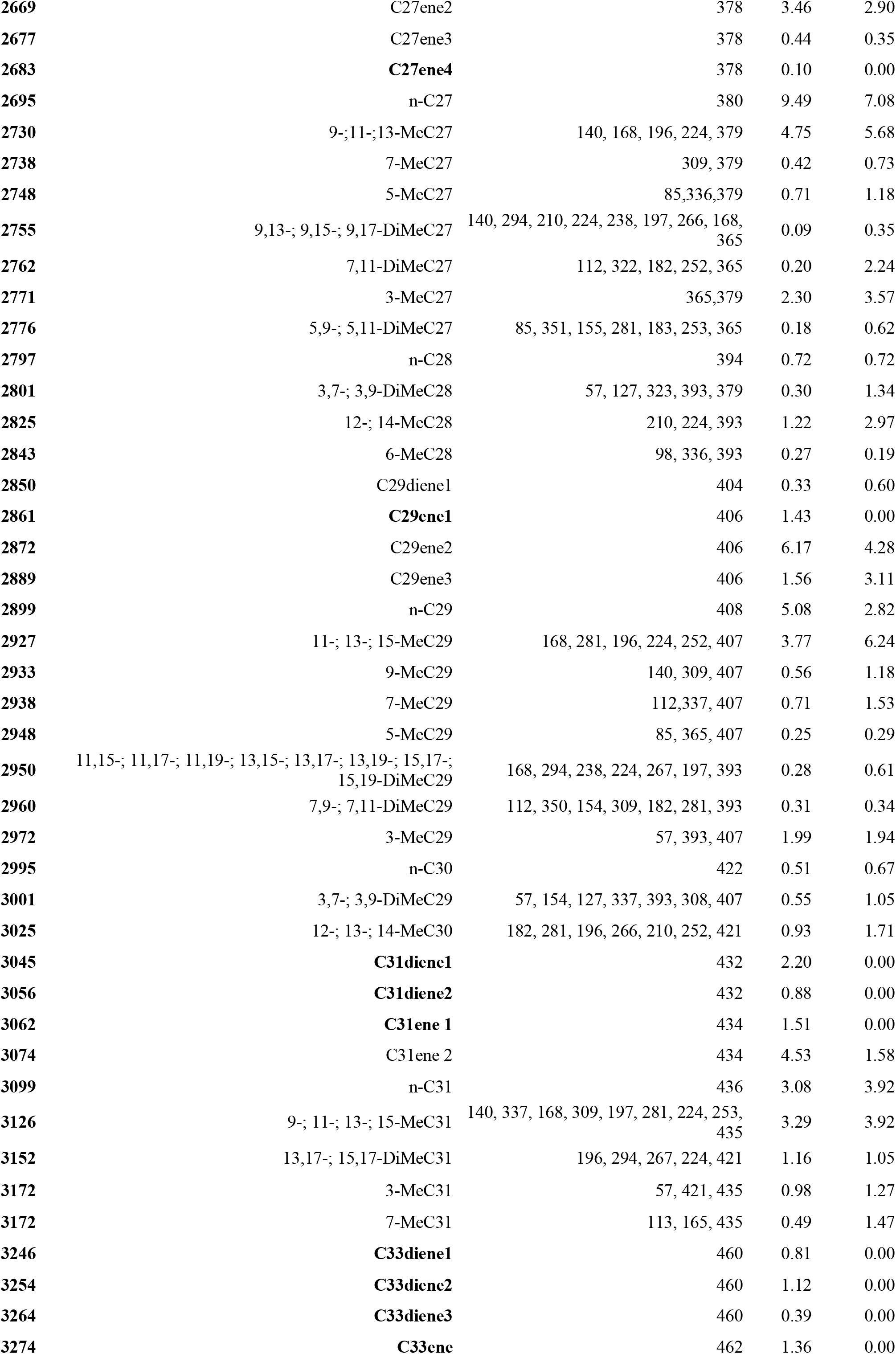
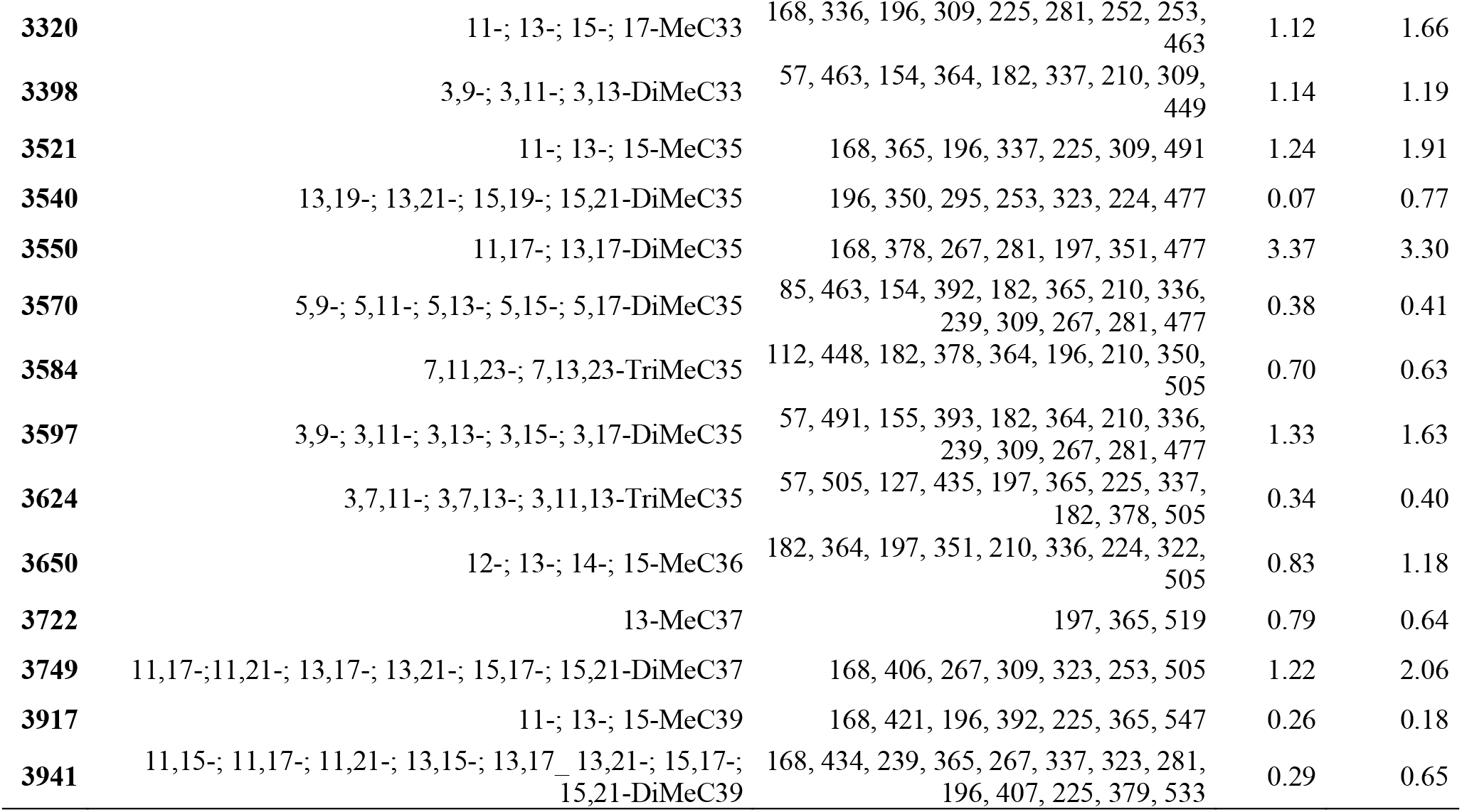
Identified cuticular hydrocarbon (CHC) compounds of alate queens and workers from Brazilian *Cardiocondyla obscurior* colonies. Indicated are the retention indices (RI), compound identifications or possible configurations in case of ambiguities, diagnostic ions obtained from the respective mass spectra, and average relative amounts per worker and queen (obtained from pools of ten individuals, respectively). Queen exclusive compounds are highlighted in bold.

| RI | Compound Name | Diagnostic Ions | queen (%) | worker (%) |
| --- | --- | --- | --- | --- |
| 1900 | n-C19 | 268 | 0.13 | 2.94 |
| 2268 | C23ene | 322 | 0.55 | 0.02 |
| 2295 | n-C23 | 324 | 0.13 | 0.17 |
| 2330 | 11-;13-MeC23 | 168,196, 323 | 0.12 | 0.10 |
| 2395 | n-C24 | 338 | 0.21 | 0.20 |
| 2468 | C25ene | 350 | 0.61 | 0.28 |
| 2499 | n-C25 | 352 | 7.26 | 4.82 |
| 2531 | 11-; 13-MeC25 | 196, 224, 351 | 2.55 | 2.07 |
| 2571 | 3-MeC25 | 337, 351 | 1.93 | 2.35 |
| 2598 | n-C26 | 366 | 1.27 | 0.93 |
| 2627 | 12-;13-MeC26 | 182, 224, 196,210, 365 | 0.69 | 1.02 |
| 2651 | 4-MeC26 | 71,337, 365 | 0.17 | 0.45 |
| 2661 | **C27ene1** | 378 | 0.13 | 0.00 |
| 2667 | C27diene | 376 | 0.29 | 0.35 |
| 2669 | C27ene2 | 378 | 3.46 | 2.90 |
| 2677 | C27ene3 | 378 | 0.44 | 0.35 |
| 2683 | **C27ene4** | 378 | 0.10 | 0.00 |
| 2695 | n-C27 | 380 | 9.49 | 7.08 |
| 2730 | 9-,11-,13-MeC27 | 140, 168, 196, 224, 379 | 4.75 | 5.68 |
| 2738 | 7-MeC27 | 309, 379 | 0.42 | 0.73 |
| 2748 | 5-MeC27 | 85,336,379 | 0.71 | 1.18 |
| 2755 | 9,13-, 9,15-, 9,17-DiMeC27 | 140, 294, 210, 224, 238, 197, 266, 168, 365 | 0.09 | 0.35 |
| 2762 | 7,11-DiMeC27 | 112, 322, 182, 252, 365 | 0.20 | 2.24 |
| 2771 | 3-MeC27 | 365,379 | 2.30 | 3.57 |
| 2776 | 5,9-, 5,11-DiMeC27 | 85, 351, 155, 281, 183, 253, 365 | 0.18 | 0.62 |
| 2797 | n-C28 | 394 | 0.72 | 0.72 |
| 2801 | 3,7-, 3,9-DiMeC28 | 57, 127, 323, 393, 379 | 0.30 | 1.34 |
| 2825 | 12-, 14-MeC28 | 210, 224, 393 | 1.22 | 2.97 |
| 2843 | 6-MeC28 | 98, 336, 393 | 0.27 | 0.19 |
| 2850 | C29diene1 | 404 | 0.33 | 0.60 |
| 2861 | **C29ene1** | 406 | 1.43 | 0.00 |
| 2872 | C29ene2 | 406 | 6.17 | 4.28 |
| 2889 | C29ene3 | 406 | 1.56 | 3.11 |
| 2899 | n-C29 | 408 | 5.08 | 2.82 |
| 2927 | 11-, 13-, 15-MeC29 | 168, 281, 196, 224, 252, 407 | 3.77 | 6.24 |
| 2933 | 9-MeC29 | 140, 309, 407 | 0.56 | 1.18 |
| 2938 | 7-MeC29 | 112,337, 407 | 0.71 | 1.53 |
| 2948 | 5-MeC29 | 85, 365, 407 | 0.25 | 0.29 |
| 2950 | 11,15-, 11,17-, 11,19-, 13,15-, 13,17-, 13,19-, 15,17-, 15,19-DiMeC29 | 168, 294, 238, 224, 267, 197, 393 | 0.28 | 0.61 |
| 2960 | 7,9-, 7,11-DiMeC29 | 112, 350, 154, 309, 182, 281, 393 | 0.31 | 0.34 |
| 2972 | 3-MeC29 | 57, 393, 407 | 1.99 | 1.94 |
| 2995 | n-C30 | 422 | 0.51 | 0.67 |
| 3001 | 3,7-, 3,9-DiMeC29 | 57, 154, 127, 337, 393, 308, 407 | 0.55 | 1.05 |
| 3025 | 12-, 13-, 14-MeC30 | 182, 281, 196, 266, 210, 252, 421 | 0.93 | 1.71 |
| 3045 | **C31diene1** | 432 | 2.20 | 0.00 |
| 3056 | **C31diene2** | 432 | 0.88 | 0.00 |
| 3062 | **C31ene 1** | 434 | 1.51 | 0.00 |
| 3074 | C31ene 2 | 434 | 4.53 | 1.58 |
| 3099 | n-C31 | 436 | 3.08 | 3.92 |
| 3126 | 9-, 11-, 13-, 15-MeC31 | 140, 337, 168, 309, 197, 281, 224, 253, 435 | 3.29 | 3.92 |
| 3152 | 13,17-, 15,17-DiMeC31 | 196, 294, 267, 224, 421 | 1.16 | 1.05 |
| 3172 | 3-MeC31 | 57, 421, 435 | 0.98 | 1.27 |
| 3172 | 7-MeC31 | 113, 165, 435 | 0.49 | 1.47 |
| 3246 | **C33diene1** | 460 | 0.81 | 0.00 |
| 3254 | **C33diene2** | 460 | 1.12 | 0.00 |
| 3264 | **C33diene3** | 460 | 0.39 | 0.00 |
| 3274 | **C33ene** | 462 | 1.36 | 0.00 |
| 3320 | 11-; 13-; 15-; 17-MeC33 | 168, 336, 196, 309, 225, 281, 252, 253, 463 | 1.12 | 1.66 |
| 3398 | 3,9-; 3,11-; 3,13-DiMeC33 | 57, 463, 154, 364, 182, 337, 210, 309, 449 | 1.14 | 1.19 |
| 3521 | 11-; 13-; 15-MeC35 | 168, 365, 196, 337, 225, 309, 491 | 1.24 | 1.91 |
| 3540 | 13,19-; 13,21-; 15,19-; 15,21-DiMeC35 | 196, 350, 295, 253, 323, 224, 477 | 0.07 | 0.77 |
| 3550 | 11,17-; 13,17-DiMeC35 | 168, 378, 267, 281, 197, 351, 477 | 3.37 | 3.30 |
| 3570 | 5,9-; 5,11-; 5,13-; 5,15-; 5,17-DiMeC35 | 85, 463, 154, 392, 182, 365, 210, 336, 239, 309, 267, 281, 477 | 0.38 | 0.41 |
| 3584 | 7,11,23-; 7,13,23-TriMeC35 | 112, 448, 182, 378, 364, 196, 210, 350, 505 | 0.70 | 0.63 |
| 3597 | 3,9-; 3,11-; 3,13-; 3,15-; 3,17-DiMeC35 | 57, 491, 155, 393, 182, 364, 210, 336, 239, 309, 267, 281, 477 | 1.33 | 1.63 |
| 3624 | 3,7,11-; 3,7,13-; 3,11,13-TriMeC35 | 57, 505, 127, 435, 197, 365, 225, 337, 182, 378, 505 | 0.34 | 0.40 |
| 3650 | 12-; 13-; 14-; 15-MeC36 | 182, 364, 197, 351, 210, 336, 224, 322, 505 | 0.83 | 1.18 |
| 3722 | 13-MeC37 | 197, 365, 519 | 0.79 | 0.64 |
| 3749 | 11,17-;11,21-; 13,17-; 13,21-; 15,17-; 15,21-DiMeC37 | 168, 406, 267, 309, 323, 253, 505 | 1.22 | 2.06 |
| 3917 | 11-; 13-; 15-MeC39 | 168, 421, 196, 392, 225, 365, 547 | 0.26 | 0.18 |
| 3941 | 11,15-; 11,17-; 11,21-; 13,15-; 13,17-; 13,21-; 15,17-; 15,21-DiMeC39 | 168, 434, 239, 365, 267, 337, 323, 281, 196, 407, 225, 379, 533 | 0.29 | 0.65 |

Overall, we identified CHCs from six different compound classes: *n*-alkanes (14.1% in queens, 16.4% in workers), *n*-alkenes (16.9% in queens, 11.5% in workers), mono-methyl-branched alkanes (35.2% in queens, 41% in workers), di-methyl-branched alkanes (21.1% in queens, 24.6% in workers), trimethylbranched alkanes (2.8% in queens, 3.3% in workers) and alkadienes (9.8% in queens, 3.3% in workers).

Comparing quantitative differences in CHC profiles of workers and alate queens, we found that alate queens have more *n*-alkenes and alkadienes than workers, but less methyl-branched alkanes (**Figure 1.A**). A principal component analysis (PCA) confirmed that CHC profiles cluster according to caste (**Figure 1.B**), separating queens and workers along principal component (PC) 1, which explains 34.1 % of the variation. The analysis further revealed that CHC variation was higher in queens than in workers, as indicated by the spread of samples from each caste. These differences were significant for PC2, but not PC1 according to Fligner-Kileen tests of equal variance (p_PC1_ = 0.127, Z_PC1_ = 1.526; p_PC2_ = 0.035, Z_PC2_ = 2.109).

**Fig. 1.**
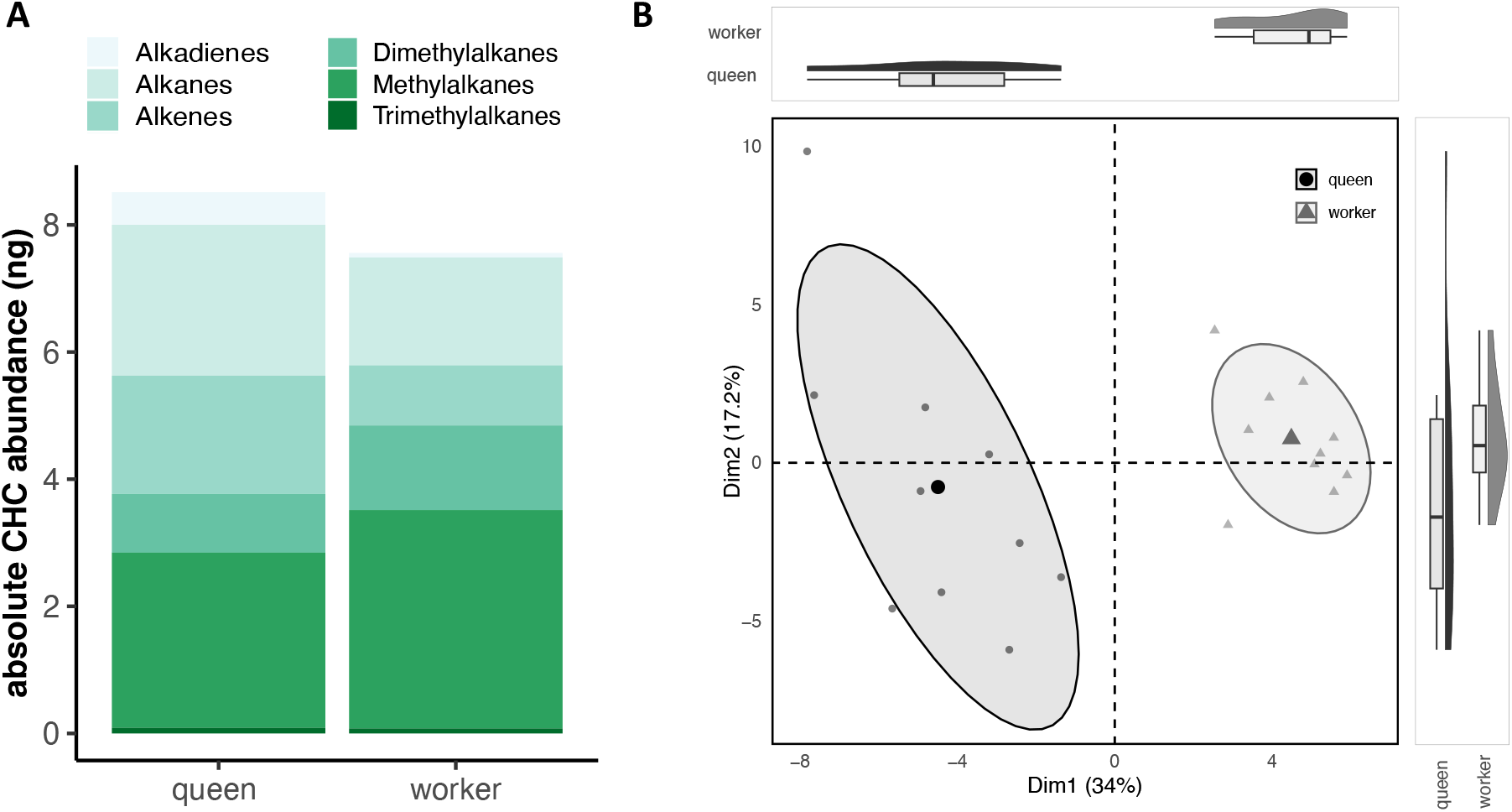
Comparison of CHC profiles in queens and workers of *Cardiocondyla obscurior*. **A** Absolute abundances (in ng) of CHC compound classes in the tested queens and workers, averaged across ten replicates each. **B** Principle Component Analysis (PCA) of CHC profiles from queens and workers of *C. obscurior*. Samples are divided by caste along Principle Component (PC) 1 (displayed on the x-axis), which explains 34.1% of the variation in the data. Additionally, compared to the worker caste, queen samples are much more dispersed on PC 2, which explains 17.4% of the variation.

To investigate how CHCs contribute to nestmate recognition and discrimination against trespassing foreign queens in *C. obscurior*, we quantified aggressive responses of workers against introduced alive, freeze-killed, and hexane-washed alate queens introduced from other nests.

At first contact, introduced foreign alate queens were always antennated by a resident worker (aggression score 1, see Box 1 for a detailed description of the typical worker response). In 31 % of the trials, no further behavioral responses were observed. In the remaining 69 % of the trials, escalating aggressive responses in which workers either showed mandible flaring (29 %), biting (8 %), pulling (28 %) or stinging (5 %) were scored. Introducing hexane-washed queens to experimental nests elicited the fewest aggressive responses, with 64 % of the trials ending with aggression score 1 (antennation). Freeze killed and alive queens were however met with substantially more aggression, with only 24 % and 7 % of the trial ending at score 1, respectively.

Ordinal logistic regression analyses with Tukey’s post-hoc testing confirmed that these differences in aggressive responses were highly significant between treatments, with hexane washed queens receiving the least aggression, followed by freeze killed queens (**Table 2, Figure 2**).

**Tab. 2.**
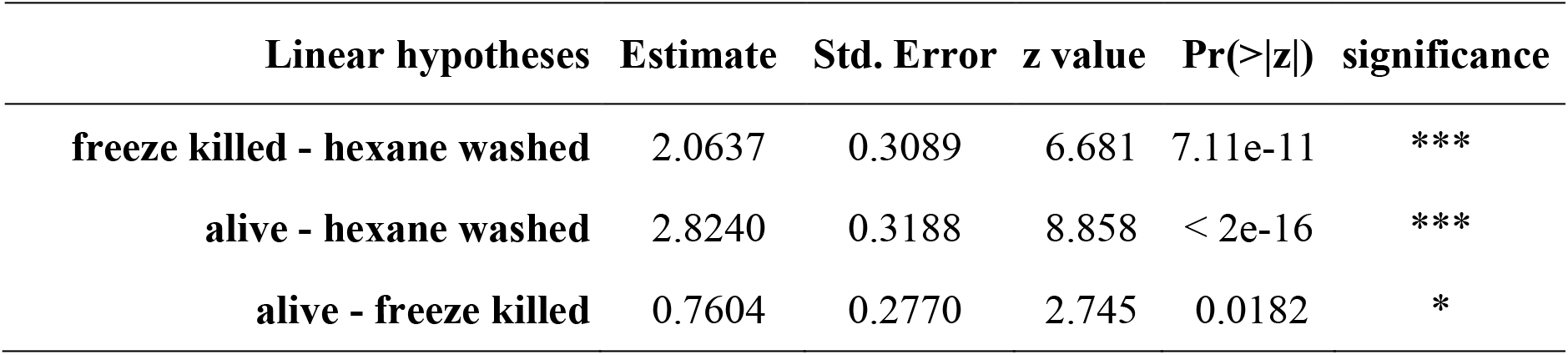
Results of Tukey’s post-hoc test (Bonferroni-corrected) of the ordinal regression analysis for aggressive worker responses against trespassing, foreign queens. Worker responses to alate queens of the respective three treatments hexane washed, freeze-killed and alive were compared, and the estimate, standard error, z value as well as Bonferroni-corrected significance levels are reported (*: p<0.05, ***: p<1e-5).

| Linear hypotheses | Estimate | Std. Error | z value | Pr(>\|z\|) | significance |
| --- | --- | --- | --- | --- | --- |
| freeze killed - hexane washed | 2.0637 | 0.3089 | 6.681 | 7.11e-11 | *** |
| alive - hexane washed | 2.8240 | 0.3188 | 8.858 | < 2e-16 | *** |
| alive - freeze killed | 0.7604 | 0.2770 | 2.745 | 0.0182 | * |

**Fig. 2.**
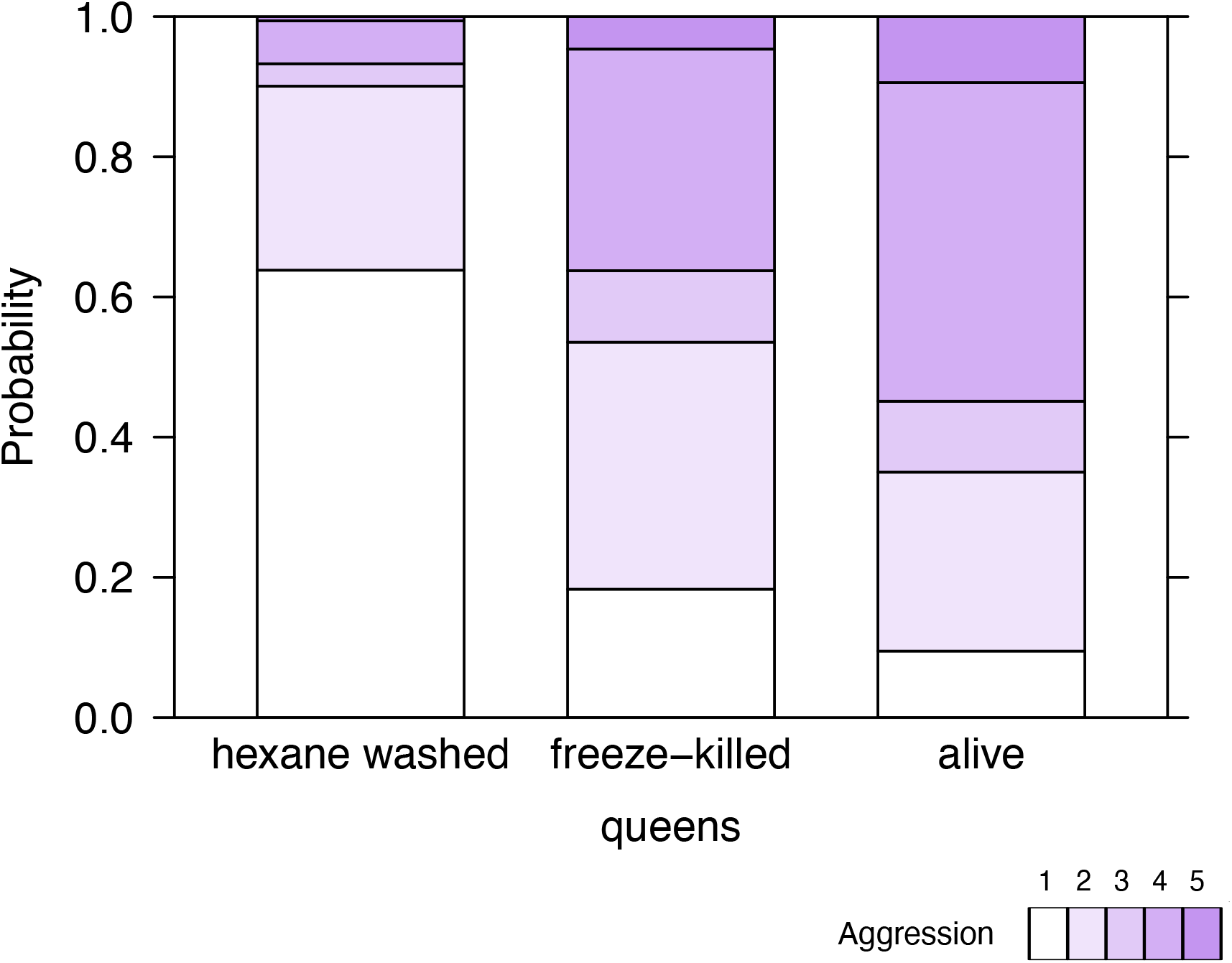
Ordinal logistic regression model of aggressive worker responses against differently treated trespassing queens of *Cardiocondyla obscurior*. The x-axis displays the different treatments of the queens (hexane washed, freeze-killed and alive) and the y-axis displays the probability of aggression scores 1 to 5. The probability to observe the lowest aggression is highest in the hexane washed treatment, whereas the probability to observe low aggression scores in alive and freeze-killed queens is significantly reduced. Aggression scores: 1= antennation, 2= mandible flare, 3=biting, 4=stinging, 5=pulling.

### Box 1

General behavior of ants in our experimental set-up

After an untreated, alive foreign queen was introduced into an experimental nest, it usually quickly encountered the resident workers. This first contact was always accompanied by antennation of all body regions (head, thorax, abdomen, wings) of the queen by one or several workers. Next, the workers either moved away or expressed escalating aggressive behaviors (mandible flaring, biting, pulling, stinging). Mandible flaring was often accompanied by intense gaster drumming. Whenever biting occurred, other workers often joined the altercation (suggesting recruitment), antennating and threatening the foreign queen before biting as well. Biting was further escalated by pulling, where one or two workers bit the queen, lifted her up, and carried her toward the nest exit. In some cases where multiple workers joined in on the aggressive behaviors, the queen was killed. Killed queens were eventually transported out of the nest.

After establishing that CHCs play a role in discrimination against trespassing queens from different colonies in *C. obscurior*, we explored whether the CHC profile differences between the antagonists (host colony profile vs. intruder colony profile) is positively correlated with the extent of aggression against an intruder.

For this, we calculated distances between CHC profiles of each colony pair based on PC1 and PC2 (which jointly explain 54.46% of the variance) to quantify the differences in CHC profiles in a single metric. To test for a significant correlation between CHC distance and aggressive response, we again used an ordinal logistic regression model with interaction of CHC distance and treatment (aggression ∼ treatment*CHC distance). This revealed that the probability for higher aggression increases with CHC distance, i.e. that aggression is positively correlated with CHC distance (**Figure 3, Table 3**). The effect was highly significant (p = 0.008) regardless of whether trespassing ants were freeze-killed or alive (i.e. no significant interaction of treatment and CHC distance, p = 0.236).

**Tab. 3.** Results of the ordinal regression analysis of aggressive worker responses with pairwise CHC distances between colonies for alive and freeze killed queens of *Cardiocondyla obscurior*. Worker responses to alate queens of the respective three treatments hexane washed, freeze-killed and alive were compared and the estimate, standard error, t value as well as significance levels are reported (*: p<0.05, **: p<0.01).

| Linear hypotheses | Estimate | Std. Error | t value | Pr(>\|z\|) | significance |
| --- | --- | --- | --- | --- | --- |
| CHC distance | 0.087 | 0.032 | 2.665 | 0.008 | ** |
| alive - freeze killed | 1.114 | 0.409 | 2.726 | 0.006 | ** |
| CHC distance - alive | -0.053 | 0.045 | -1.184 | 0.236 |  |

**Fig. 3.**
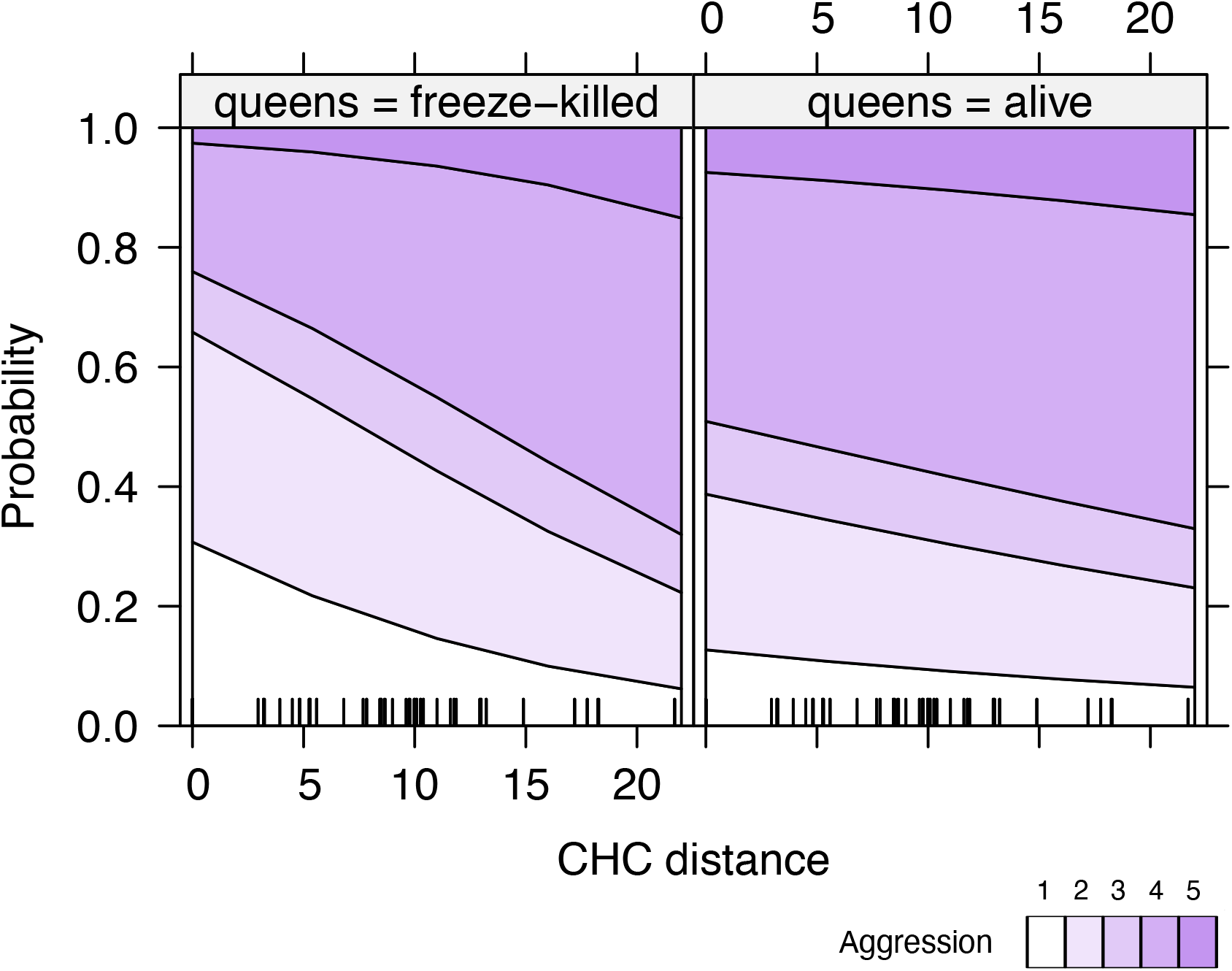
Ordinal logistic regression model of the effect of pairwise CHC distances on the aggressive worker responses against alive or freeze-killed queens of *Cardiocondyla obscurior*. Pairwise CHC distances were calculated based on the first two principal components of the alate queen CHC profiles from each colony. The x-axis displays the pairwise CHC distance and the y-axis shows the modelled probability of different aggression scores. The probability to observe higher aggression increases significantly with CHC distance in both treatments.

## Discussion

In this study, we used behavioral assays and CHC profiling of colonies of *Cardiocondyla obscurior* from an invasive population to explore mechanisms of intraspecific aggression. We describe qualitative and quantitative differences in CHC profiles of alate queens and workers and show a higher chemical diversity in queens than in workers. We also demonstrate experimentally that the cues to elicit aggression of workers against foreign, trespassing queens are encoded in their CHC profiles. Finally, we show that aggression correlates with CHC profile divergence between interacting partners, i.e. that aggression is higher the more divergent the CHC profiles are.

Together, our findings suggest that invasive populations of *C. obscurior* in Brazil are multi-rather than unicolonial. The high levels of aggression against trespassing reproductives from foreign colonies potentially impede unhindered movement of individuals among nests, thus precluding unicoloniality in these populations (HelanterÄ & al. 2009). Therefore, unicoloniality does not appear to be a necessary condition for invasive populations of *C. obscurior* to persist, consistent with results obtained for populations of two other invasive ants, *Myrmica rubra* (Garnas & al. 2007, Chen & al. 2018) and of *B. patagonicus* (Eyer & al. 2021).

We can further conclude that colonies are polydomous in these populations of *C. obscurior*, as aggression is significantly reduced between individuals with more similar CHC profiles, which is most likely to occur between colonies that are highly related and that share the same microenvironment (Menzel & al. 2017). Such networks of highly related colonies in close proximity are likely to develop in ant species such as *C. obscurior* where budding is the primary mode of colony propagation (Heinze & al. 2006, HelanterÄ & al. 2009).

Previous studies already documented antagonistic interactions between workers (Heinze & al. 2006) and increased levels of aggression against foreign queens in *C. obscurior* (Schrader & al. 2014). However, the role of CHCs in these interactions had not yet been explored. Based on our experiments we here provide first empirical evidence that CHCs play a key role in intercolonial aggression against trespassing queens in this species. Showing that CHC-depleted queens received virtually no aggression from workers compared to untreated or freeze-killed queens, we can conclude that aggression is predominantly mediated by chemical cues encoded in the CHCs.

Earlier studies on queen CHC profiles in *C. obscurior* found decidedly fewer individual compounds by exclusively analyzing thermally desorbed volatized CHCs (Will & al. 2012, Schrader & al. 2014). Here, by using solvent extractions, we were able to identify 71 compounds in queen CHC profiles, demonstrating that solvent extractions of surface profiles are better suited to provide a comprehensive representation of CHC profiles. Accordingly, our approach allowed for the identification of ten compounds exclusively found in alate queens, which have not been reported so far in this species. All these compounds are unsaturated CHCs, namely *n*-alkenes and alkadienes (Supplementary Table **S1**). Such compounds are well-known chemical communication signals in insects, most prominently in *Drosophila* where they constitute both the main male and female sex pheromonal compounds (Dallerac & al. 2000, Marcillac & Ferveur 2004, Grillet & al. 2006). Furthermore, a C29 alkene, of which we also found several in *C. obscurior* CHC profiles (albeit only one as queen exclusive), has been identified as a queen fertility signal across multiple geographically varied populations of the trap-jaw ant *Odontomachus brunneus* Patton (Smith & al. 2013, Smith & al. 2015). Therefore, the queen-specific *n*-alkenes and alkadienes identified in our analyses are promising targets for further studies to resolve why foreign alate queens are met with significantly more aggression than foreign workers in this species (Schrader & al. 2014).

Our population-level screen of CHC profiles also revealed that alate queen profiles in general are more variable compared to those of workers. A foreign alate queen might thus be recognized and discriminated against more easily by a colony, as her CHC profile is more likely to be distinct from the profiles of nestmate queens. In contrast, CHC profiles of workers were found to be more uniform, which likely reduces the ability of a colony to discriminate between nestmate and non-nestmate workers as opposed to queens. Heinze *et al*. (2006) observed that foreign workers are eventually integrated into a colony after being met with only low levels of aggression. If adoption is common, then the adaptive benefit of recognizing and discriminating against foreign reproductives is obvious, as adoption of foreign workers (which are entirely sterile in *Cardiocondyla*) is unlikely to reduce the reproductive output of a colony (Smith & Loope 2016). If anything, it is more likely to increase it, as workers will contribute to the communal resources of the colony (Kennedy & al. 2021). Adoption of foreign queens, however, is much more likely to reduce the fitness of an individual colony, as a queen will consume a colony’s resources to maximize her own reproductive output.

Populations of invasive ants commonly show low genetic and chemical diversity due to genetic bottlenecks, which is expected to result in reduced aggressive behavior (Tsutsui & al. 2000, Tsutsui & al. 2003). This, in turn has been hypothesized to drive the emergence of unicolonial populations. Unicoloniality has been shown to give invasive ants a competitive advantage over native species, as the invaders can quickly reach high population densities without the negative effects of intraspecific conflicts (Heinze & al. 2006, Kennedy & al. 2021). For *C. obscurior*, Errbii *et. al*. (2021) showed that intraspecific hybridization between two divergent genetic lineages and the activity of transposable elements increase genetic diversity, which could increase the diversity at the level of CHCs and thus explain the relatively high levels of aggression. This might also explain why the ecological impact of invasive *C. obscurior* appears to be smaller than that of other invasive but supercolonial species of ants (Holway & al. 2002, Klotz & al. 2008, Siddiqui & al. 2021). This further invites detailed population genetic studies investigating the correlation of physical and genetic proximity in C. *obscurior* and their collective impact on CHC profile divergence.

Intriguingly, intraspecific aggression appears to be a polymorphic trait in *C. obscurior*. Schrader *et al*. (2014) reported much lower levels of intraspecific aggression in a genetic lineage that diverged from the Brazil population about 10,000 years ago (Errbii & al. 2021). This more docile variant of *C. obscurior* also has a much broader distribution range, with populations found e.g. in Taiwan, Japan, Spain, North America and green-houses in Europe (Errbii & al. unpubl.). Whether this apparent higher invasive potential can be explained by the reduced aggression and whether the reduced aggression can be explained by reduced chemical diversity or by other means however remains to be investigated.

Ants have evolved sophisticated means of chemical communication to precisely regulate social interactions. Our understanding of the intricate details remains only superficial, also because of the considerable variation that evolved in this system across different species, genera, and clades within the Formicidae. Here, we have taken a step towards better understanding chemical communication in *C. obscurior*. Our study provides a foundation for future work in this species to decipher the chemical signals underlying antagonistic interactions, the associated behavioral polymorphism, and the consequences for the species’ invasive potential.

## Acknowledgments

We thank Tobias van Elst for help with collecting ants in the field. This research was funded by the Deutsche Forschungsgemeinschaft (DFG, German Research Foundation) – 403813881 with a grant to L.S. (SCHR 1554/2-1) under the priority program “Rapid evolutionary adaptation: Potential and constraints” (SPP 1819). We are grateful to Jacques Delabie for supporting field work in Brazil and the Brazilian Ministério do Meio Ambiente for permission to study the introduced populations of *Cardiocondyla obscurior* in Bahia (permit 63371-1).

