## Supplementary Figure S1 for "The role of cuticular hydrocarbons in intraspecific aggression in the invasive ant *Cardiocondyla obscurior*"

**Supplementary Material for "The role of cuticular hydrocarbons in intraspecific aggression in the invasive ant *Cardiocondyla obscurior*"**


Maja DRAKULA, Jan BUELLESBACH, Lukas SCHRADER

**Address(es) of author(s)**

Lukas Schrader (contact author), Institute for Evolution and Biodiversity, University of Münster, Hüfferstraße 1, 48149 Münster, Germany.


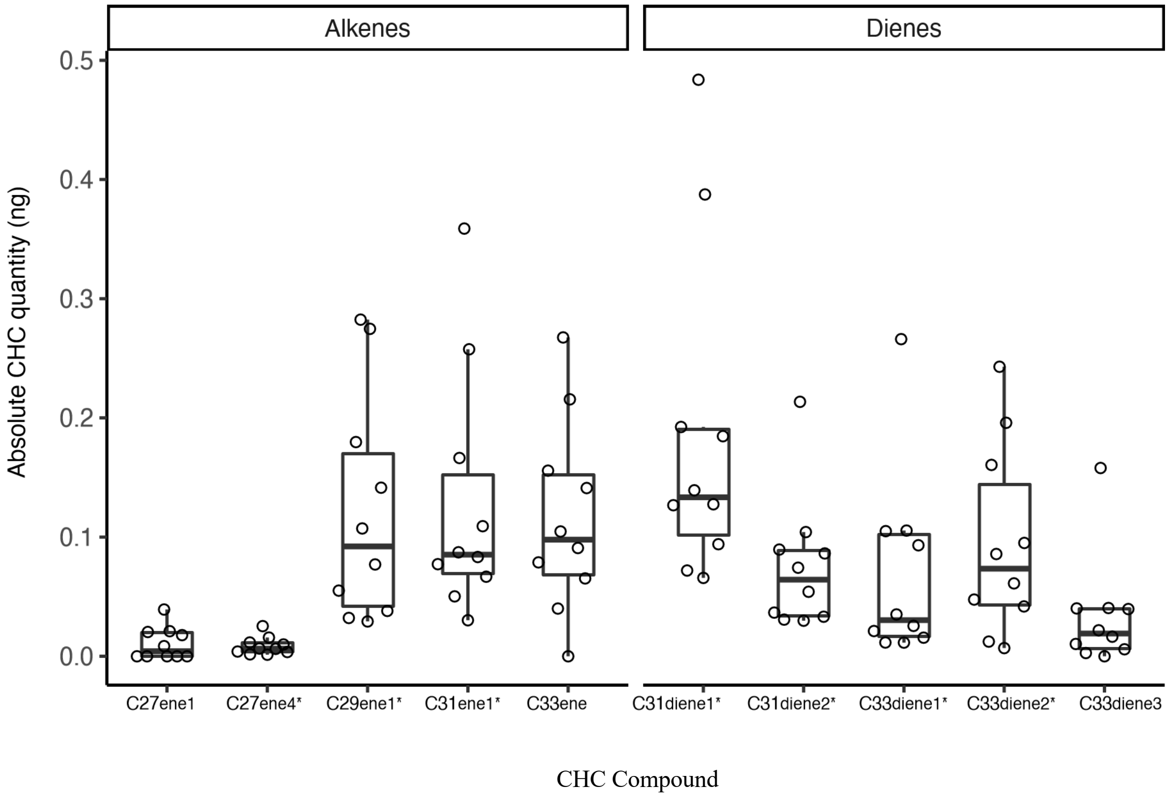


**Supplementary Figure S1: Absolute quantity of the ten CHC compounds identified in alate queens but not workers of *Cardiocondyla obscurior*.** Three of the five alkenes and four of the five dienes were detected in each of the ten analyzed queen samples (highlighted with an asterisk).
