## Supplementary Table S1 for "The role of cuticular hydrocarbons in intraspecific aggression in the invasive ant *Cardiocondyla obscurior*"

**Supplementary Material for “The role of cuticular hydrocarbons in intraspecific aggression in the invasive ant *Cardiocondyla obscurior*”**

Maja DRAKULA, Jan BUELLESBACH, Lukas SCHRADER

**Address(es) of author(s)**

Lukas Schrader (contact author), Institute for Evolution and Biodiversity, University of Münster, Hüfferstraße 1, 48149 Münster, Germany.

**Supplementary Table S1: Colonies of *C. obscurior* used for chemical analysis and behavioral assays.** The table includes the Sample ID, GPS location of the collection area, location, as well as the total numbers of queens and workers used for chemical analysis and behavioral assays, and the respective values for the Principal Components (1 and 2) for each caste.

| Sample ID | GPS | Location | Queen CHC profiles assessed | Worker CHC profiles assessed | # queens for behavioral assays | PC1 queens | PC2 queens | PC1 workers | PC2 workers |
|---|---|---|---|---|---|---|---|---|---|
| 007 | (-15.2634637,-39.0833540) | Una, Brazil | 10 | 10 | 27 | -2.6789 | -5.8953 | 4.7953 | 2.5565 |
| 054 | (-15.2861191,-39.0627775) | Una, Brazil | 10 | 10 | 27 | -5.6537 | -4.5976 | 2.5419 | 4.1793 |
| 080 | (-15.2207770,-39.0357294) | Una, Brazil | 10 | 10 | 27 | -7.6400 | 2.1357 | 5.0796 | -0.0437 |
| 082 | (-15.2222370,-39.0358943) | Una, Brazil | 10 | 10 | 27 | -1.3626 | -3.6058 | 5.5701 | 0.7951 |
| 081 | (-15.2217323,-39.0351846) | Una, Brazil | 10 | 10 | 27 | -3.1915 | 0.2676 | 5.2294 | 0.2939 |
| 013 | (-14.7914687,-39.1872256) | Itabuna, Brazil | 10 | 10 | 27 | -4.9344 | -0.8904 | 5.9074 | -0.3913 |
| 014 | (-14.7914464,-39.1872531) | Itabuna, Brazil | 10 | 10 | 27 | -4.8145 | 1.7505 | 3.4024 | 1.0415 |
| 015 | (-14.7623629,-39.2327735) | Itabuna, Brazil | 10 | 10 | 27 | -4.4035 | -4.0826 | 2.8816 | -1.9577 |
| 025 | (-14.7916513,-39.1866942) | Itabuna, Brazil | 10 | 10 | 27 | -7.8180 | 9.8260 | 5.5734 | -0.9130 |
| 024 | (-14.7917329,-39.1865088) | Itabuna, Brazil | 10 | 10 | 27 | -2.4150 | -2.5309 | 3.9309 | 2.0621 |
